# Zebrafish (*Danio rerio*) behavioral laterality predicts increased short-term avoidance memory but not stress-reactivity responses

**DOI:** 10.1101/565309

**Authors:** Barbara D. Fontana, Madeleine Cleal, James M. Clay, Matthew O. Parker

**Author notes:** Correspondence to: **Matthew O. Parker, PhD**, School of Pharmacy and Biomedical Sciences, University of Portsmouth, Old St Michael’s Building, White Swan Road, Portsmouth, PO1 2DT, UK., **Barbara D. Fontana**, School of Pharmacy and Biomedical Sciences, University of Portsmouth, Old St Michael’s Building, White Swan Road, Portsmouth, PO1 2DT, UK.

## Abstract

Once considered a uniquely human attribute, behavioral laterality has proven to be ubiquitous among non□human animals, being frequently associated with different neurophenotypes in rodents and fish species. Zebrafish (*Danio rerio*) are a **versatile vertebrate model system** that has been widely used in translational neuropsychiatric research due their highly conserved genetic homology, well characterized physiological and extensive behavioral repertoire. Although the spontaneous left- and right-bias responses and associated behavioral domains (e.g. stress reactivity, aggression and learning) have previously been observed in other teleost species, no information regarding **how spontaneous motor left-right bias responses of zebrafish predicts other behavioral domains** has been described. Thus, we aimed to investigate the existence and incidence of natural left-right bias of adult zebrafish in the Y-maze test and explore any relationship of biasedness on the performance of different behavioral domains. This included learning about threat-cues in the fear conditioning test and locomotion and anxiety-related behavior in the novel tank diving test. In conclusion, we showed for the first time that zebrafish exhibit a natural manifestation of **motor** behavioral lateralization which can influence aversive learning responses. **Although laterality did not change locomotion or anxiety-related behaviors, we found that biased animals had an altered exploration pattern in the Y-maze, making them easily discernable from their unbiased counterparts, and increased learning associated to fear cues.**

## 1. Introduction

Lateralization of brain and behavior is the apparent predisposition towards side bias, often manifested in terms of motor output, such as handedness. Laterality has been extensively studied in humans, **with functional laterality of different brain regions known to be an important aspect in language and cognition (Hull and Vaid 2007; Vikingstad et al. 2000). Research has revealed that motor functions may also be under the control of lateralized mechanisms which, in humans, may manifest as preference for one side over the other for handedness, footedness and eyedness** (Brown and Taylor 1988). **Laterality** has been shown to be an evolutionarily conserved characteristic which is observed at the populational level and, when disrupted, has been associated with **cognitive and neuropsychiatric disorders, such as anxiety and major depressive disorder** (Koster et al. 2010; Lichtenstein-Vidne et al. 2017).

**Lateralization is not a purely human characteristic, many studies now suggest that most vertebrate species**, including monkeys (Fagot and Vauclair 1991; Hopkins 1994; McGrew and Marchant 1997), rodents (Robison 1981; Rodriguez and Afonso 1993; Rodriguez et al. 1992), birds (Bhagavatula et al. 2014; Franklin and Adams 2010; Gunturkun et al. 1998), fish (Bibost and Brown 2014; Bisazza and de Santi 2003; Dadda et al. 2010a; Dadda et al. 2010b) **and some invertebrate species (Anfora et al. 2010, 2011; Frasnelli, Vallortigara, and Rogers 2012) express brain functional asymmetries (MacNeilage, Rogers, and Vallortigara 2009). In rodents for example**, several behavioral tasks have been used to assess behavioral asymmetries such **as** turning rotometers, handedness, choice behavior, T-maze and Y-maze (Corballis 1986; Pisa and Szechtman 1986; Zimmerberg and Glick 1974). Variability in lateralization exerts a number of fitness benefits **at the individual level**. For example, lateralization has been associated with maximization of brain processes, enabling individuals to process two tasks simultaneously (Rogers 2000; Rogers 2002). **Studies have suggested that laterality evolved at the population level to maintain coordination among social groups once that lateralization in a same direction is considered as a potential benefit to group cohesion and a mechanism where individuals interact in predictable ways in social group** (Rogers 2000).

Behavioral laterality is an evolutionarily conserved characteristic which is observed at populational level in humans and has been associated to different neurophenotypes (Corballis 2017; Frasnelli 2013). **Taxonomic and evolutionary aspects of brain laterality have been previously described in fish species, primarily focusing on CNS asymmetries, sensory organs and somatic lateralization, as well as the adaptive role of laterality in nature (de Perera and Braithwaite 2005; Lychakov 2013; Nepomnyashchikh and Izvekov 2006; Vallortigara and Rogers 2005). Behavioral** asymmetries have been related to high escape performance (Dadda et al. 2010b), social responses (Reddon and Balshine 2010) and even accelerated learning responses (Andrade et al. 2001) in **both fish and mammals. However, despite the clear** relevance of lateralization to several mammals and vertebrate species, we still have a limited understanding of the general origins of morphological and functional asymmetries in the brain and of their importance for behavior.

Zebrafish (*Danio rerio*) is a versatile vertebrate model system that has been widely used in translational neuropsychiatric research (Fontana et al. 2018; Stewart et al. 2015) and **to understand evolutionary aspects of animal cognition** (Oliveira 2013; Reale et al. 2007). **The last decade has seen an increase in the use of zebrafish to study the mechanisms underlying lateralization** (Andersson et al. 2015; Ariyomo and Watt 2013; Barth et al. 2005; Dadda et al. 2010a; Sovrano and Andrew 2006). **Larval zebrafish have an increased use of left-eye when interacting with their own reflection** (Sovrano and Andrew 2006). **Studies also showed that zebrafish initially use the right hemifield predominantly when interacting with novel object, but as the object becomes familiar, they switch to the left hemifield** (Miklósi, Andrew, and Savage 1997). **In a mutant zebrafish strain that shows strong laterality bias (*fsi*), asymmetry of diencephalic structures correlates with different behavior patterns such as boldness responses, with left-bias fish showing increased latency to interact with a novel object than right-bias fish** (Barth et al. 2005). **Despite a growing understanding of lateralization in zebrafish, very little is known about how spontaneously occurring motor asymmetry impacts on other behavioral domains. Therefore, in this study we aimed: (1) to investigate the existence and incidence of spontaneously occurring motor left-right bias of adult zebrafish in a continuous Y-maze test; and (2) to explore how spontaneously occurring motor left-right bias relates to performance on different behavioral domains, including learning about threat-cues in the fear conditioning test, and locomotion and anxiety-related behavior in the novel tank test.**

## 2. Material and Methods

### 2.1. Animals

**Adult zebrafish (AB wild-type; ~ 50:50 male:female ratio at 3-month of age) were bred in-house and reared in standard laboratory conditions on a re-circulating system** (Aquaneering, USA). Animals were maintained on a 14/10-hour light/dark cycle (lights on at 9:00 a.m.), pH 8.4, at ∼28.5 °C (±1 °C) in groups of 20 animals per 2.8 L. Fish were fed three times/day with a mixture of live brine shrimp and flake food, **except on the weekends when they were fed once a day**. **Animals were tested in the Y-maze apparatus and then pair-housed for 24 hours prior to analysis of shock-avoidance or tank diving**, to reduce stress from multiple handling in a single day (see Fig. 1). **During the pair-housing period, animals had shared water system and visual contact through transparent partition, the pair-housing system is used to separate animals for further identification and reduce the stress induced by social isolation (Parker et al. 2012).** After behavioral tests, all animals were euthanized using 2-phenoxyethanol from Aqua-Sed (Aqua-Sed™, Vetark,Winchester, UK). All experiments were carried out following approval from the University of Portsmouth Animal Welfare and Ethical Review Board, and under license from the UK Home Office (Animals (Scientific Procedures) Act, 1986) [PPL: P9D87106F].

**Figure 1.**
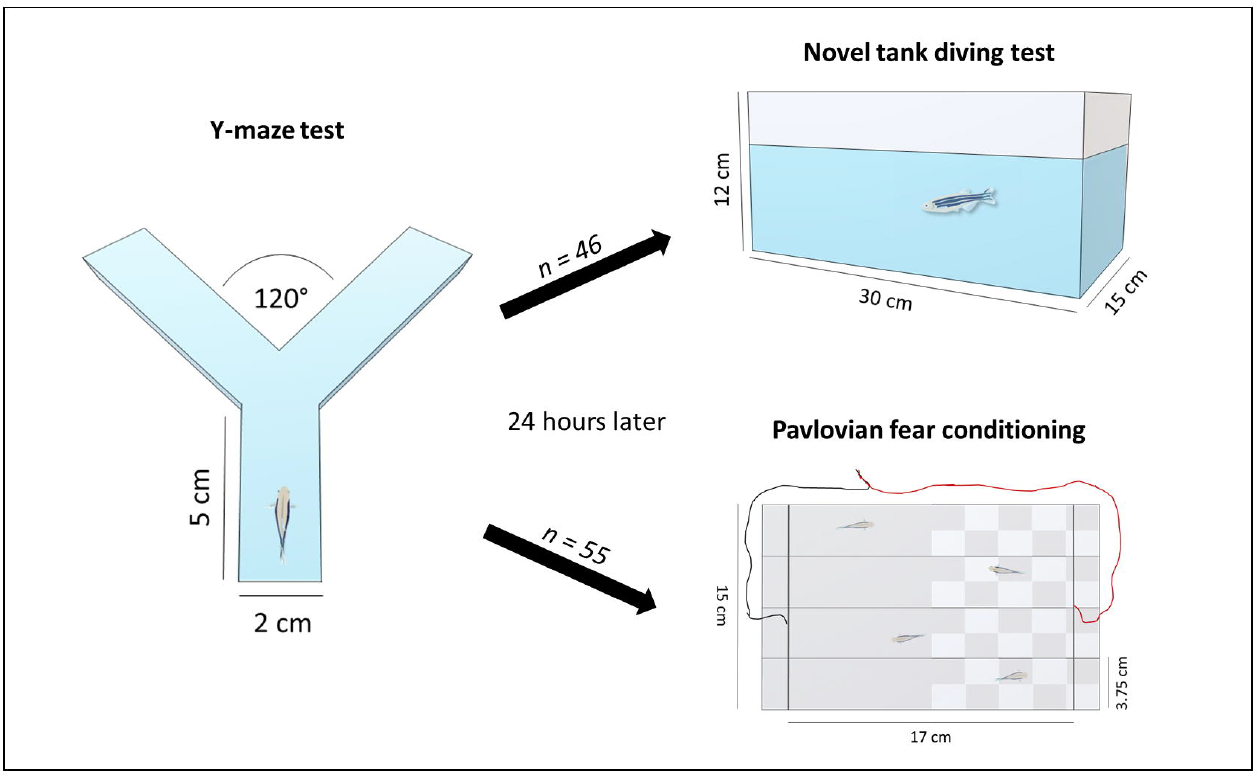
Schematic representation of the experimental design and the behavioral tasks.

### 2.2. Y-maze test

The Y-maze spontaneous alternation task is widely used for measuring the disposition of different animal models to explore new environments and to assess left- and right-biased responses (Barnard et al. 2016; Castellano et al. 1987; Frasnelli 2013; Rodriguez et al. 1992). **One-hundred and three** adult zebrafish were used for assessing Y-maze performance and right-left bias. Required sample size was calculated *a priori* following pilot tests (effect size (d) = 0.3, power = 0.8, alpha = 0.05). The apparatus consisted of a white Y-maze tank with three identical arms (5 cm length x 2 cm width; three identical arms at a 120º angle from each other) and a transparent base, filled with 3L of aquarium water (Fig. 1). Ambient light allowed some visibility in the maze, but no explicit intra-maze cues were added to the environment. **Behavioral tests were performed between 10 a.m. to 4 p.m. using the automated Zantiks (Zantiks Ltd., Cambridge, UK) AD system (Brock et al. 2017) to track animal behavior, and we carried out three independent replicate**s. **The Zantiks AD system was fully controlled via a web-enabled device during behavioral training. Fish behavior patterns were recorded for 1 hour** and was assessed according to overlapping series of four choices (tetragrams) and analyzed as a proportion of the total number of turns (Gross et al. 2011). **For assessing fish behavioral lateralization, the number of turns for the right or left arm according to the previous location in 10 minute time bins was used and the following formula was applied:** 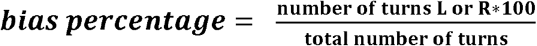. Following this, the mean and coefficient of variation for right and left bias was calculated based on the bias percentage in 10-minute time bins to assess the consistency of data across the entire trial (1-hour test period). Behavioral lateralization was considered when the animal presented >60% of average preference for right or left turns in the Y-maze during the 1-hour test. To assess zebrafish search strategies we used the number of alternations (rlrl + lrlr) and repetitions (rrrr + llll) as proportion of total number of turns which are highly expressed through 1-hour and are commonly linked to animals’ exploratory activity (Cleal and Parker 2018; Gross et al. 2011). Both alternations and repetitions were also previously linked to short-term spatial memory by reduction the number of alternations after developmental alcohol exposure (Cleal and Parker, 2018); however, more studies are need to clarify how these patterns are linked to learning and memory processes.

### 2.3. Pavlovian fear conditioning

The inhibitory avoidance paradigm is a valid method widely used to explore mechanisms underlying fear avoidance learning responses in zebrafish (Amorim et al. 2017; Manuel et al. 2014; Manuel et al. 2015; Ng et al. 2012). 24h after the completion of the Y-maze test, animals (*n*= 55) were tested on a Pavlovian fear conditioning procedure for 1 hour. The fear conditioning response was based on previous work (Cleal and Parker 2018; Valente et al. 2012) **where increased memory responses were linked to the decreased time spent in the fear conditioned area**. Fish were individually placed in one of four lanes of a tank (25 cm length x 15 cm, 1 L of water in each tank) **of the Zantiks (Zantiks Ltd., Cambridge, UK) AD system** (Brock et al. 2017) (Fig. 1). Briefly, fish were habituated for 30 minutes in the test environment**, half of the arena had a check pattern on the base and the other half had a grey color on the base. Images were projected into the tank from a screen which was embedded in the base of the AD unit, on which the test tank sits. Every 5 minutes of the habituation the base patterns, check or grey, were switched to the opposite side of the tank. The baseline preference was ascertained over 30-mins, but only the last 10-mins baseline preference was used for assessing the area preference.** Following the baseline was the conditioning phase, which consisted of a conditioned stimulus (**CS+; full screen of “check” or “grey”, half CS+ in “check” and CS+ in “grey”**) presented for 1.5-s and followed by a brief mild shock (9 V DC, 80ms; unconditioned stimulus (US). After this, an 8.5-s inter-trial interval (ITI) of the non-CS (CS−) exemplar was presented at the bottom of the tank. The CS+/US was **shown** nine times. Finally, avoidance of CS+ was assessed by presenting the baseline screen (CS+ and CS− simultaneously) for 1-min, and switching positions after 30-s.

### 2.4. Novel tank diving and thigmotaxis responses

The novel tank test is commonly used for analyzing locomotor and anxiety-like phenotypes in zebrafish presenting a high sensitivity to anxiolytic and anxiogenic drugs (Egan et al. 2009; Kalueff et al. 2013; Levin et al. 2007; Maximino et al. 2010; Mezzomo et al. 2016; Wong et al. 2010). 24 hours after the Y-maze test, animals (*n = 46*) were placed individually in a novel tank (30 cm length x 15 cm height x 12 cm width) containing 4 L of aquarium water. Behavioral activity was recorded using 2 webcams (front and top view, see Fig. 1B) for 5 min to analyze thigmotaxis and diving response (Egan et al. 2009; Parker et al. 2012; Rosemberg et al. 2012). Behaviors were **measured using an automated** video-tracking software (EthoVision, Noldus Information Technology Inc., Leesburg, VA - USA) at a rate of 60 frames/s. **Thigmotaxis (time spent in proximity to the edge/sides) was analyzed through the behavior tracking obtained by the top view camera. Meanwhile, for the detailed evaluation of vertical activity, the tank was separated in three virtual areas (bottom, middle and top) and the** following endpoints were measured: total distance traveled, time spent in each third of the tank, immobility and thigmotaxis.

### 2.5. Randomization and blinding

All behavioral testing was carried out in a fully randomized order, choosing fish at random from one of ten housing tanks for testing. Fish were screened for left-right bias in the Y-maze first, but analysis was not carried out prior to subsequent behavioral testing to avoid bias. Subsequent to **the** Y-maze screening, fish were pair housed and issued a subject ID, allowing all testing to be carried out in a fully blinded manner. **Once all data were collected and screened for extreme outliers (e.g., fish freezing and returning values of ‘0’ for behavioral parameters indicating non-engagement), the bias was revealed, and data analyzed in full. Two animals were excluded in the Y-maze task and two in the novel tank due to poor engagement with the task (freezing and displaying no measurable behavioral patterns)**.

### 2.6. Data processing and statistical analysis

**Data from the Y-maze protocol was obtained as number of entries into each arm (1, 2, 3 and middle section 4) across a 1-hour trial. To analyze the data according to left and right turns in 10-minute time bins, raw data was processed using the Zantiks Y-maze Analysis Script created specifically for this purpose (available from: https://github.com/thejamesclay/ZANTIKS_YMaze_Analysis_Script).** Subsequently, data were analyzed in IBM SPSS® Statistics and the results were expressed as means ± standard error of the mean (S.E.M), to assess whether there were any effects of bias on total turns, alternations (lrlr + rlrl) **or** repetitions (rrrr + llll). **Alternations and repetitions were analyzed using generalized linear models (Poisson distribution, log link), with laterality (three levels – left bias, right bias, non-bias) and time (six levels - 10-min time bins across 1 hour) as the fixed factor, and ID as a random effect (to account for non-independence of replicates). To analyze novel tank responses, we used either one- or two-way ANOVAs, with bias and time spent in different tank zones fixed factors (tank diving) or bias only as a fixed factor for thigmotaxis, immobility and distance traveled.** Additionally, left-right bias effects on **the** shock avoidance test was assessed using two-way repeated measures ANOVA with ‘bias’ (left vs right vs neutral) and conditioning (pre-vs-post) as factors, and preference for conditioned stimulus as the dependent variable. Newman-Keuls test was used as post-hoc analysis, and results were considered significant when p ≤ 0.05.

## 3. Results

### 2.1. Left-right bias profile in the Y-maze test

Zebrafish showed behavioral lateralization in the Y-maze (right-biased 27.18 %, left-biased 27.18% and non-biased 45.63%). To confirm if the behavioral laterality was **consistent across 1-hour (analyzed in 10-minute time bins)**, the coefficient of variation for the left and right-turn preferences were calculated for the non-biased (19.28 ± 2.52 and 21.05 ± 2.95), left-biased (30.40 ± 3.85 and 21.7 ± 3.61) and right-biased (27.23 ± 5.52 and 25.95 ± 2.31) groups. Figure 2 displays the Y-maze data. A significant bias effect was observed for number of turns (F _(2, 601)_ = 13.115; *p*<0.0001), repetitions (F _(2, 601)_ = 39.696; *p*<0.0001) and alternations (F _(2, 601)_ = 45.437; *p*<0.0001). A time effect (data not shown) was also observed for number of turns (F _(5, 601)_ = 9.769; *p*<0.0001), repetitions (F _(5, 601)_ = 3.242; *p*=0.007) and alternations (F _(5, 601)_ = 3.801; *p*=0.002). Additionally, a significant interaction effect (bias *\** time) was observed for repetitions (F _(10, 601)_ = 2.504; *p*=0.006) and alternations (F _(10, 601)_ = 2.390; *p*= 0.009). **Bias, to the left or the right, significantly increased the number of repetitions (*p*<0.0001 for right-bias and *p*<0.001 for left-bias) and decreased the percentage of alternations (*p*<0.0001 for right-bias and *p*<0.005 for left-bias) compared to non-biased animals. In addition, right-biased animals also decreased the number of turns (p<0.05) compared to non-biased animals. Biased differences were not just seen between biased vs. non-biased groups, but also between bias groups**, right-biased animals had a significant increase of repetitions (*p*<0.05) and decrease of alternations (*p*<0.05) compared to left-biased animals (Fig. 2A). The behavioral profile of biased and non-biased animals is displayed in Fig. 2B tetragrams where a high number of llll and rrrr configuration can be observed for left- and right-biased animals, respectively.

**Figure 2.**
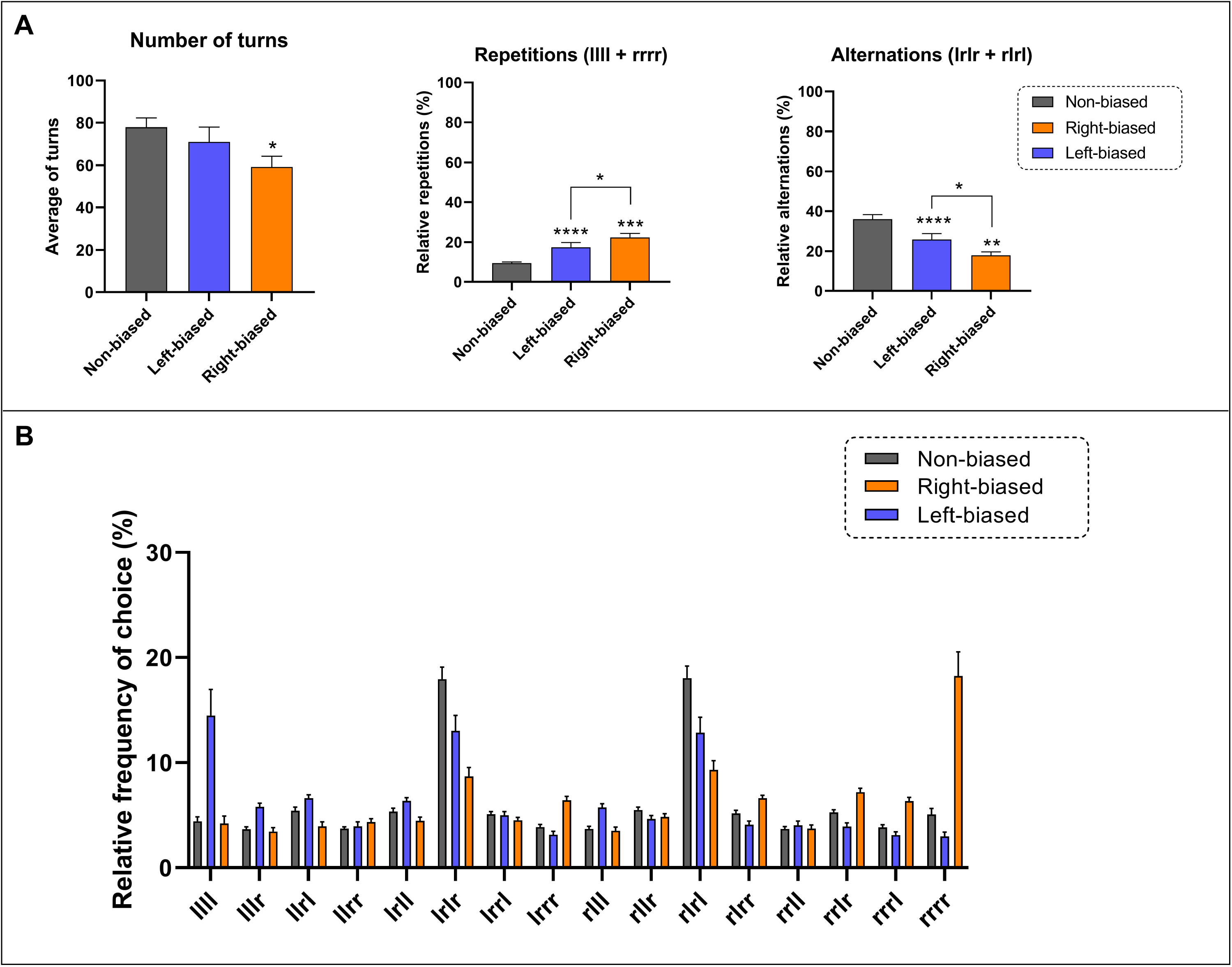
Effects of left- and right-bias in zebrafish on the Y-maze test. (A) Laterality affects total number of turns, repetitions and alternation of adult zebrafish. **(B) Y-maze tetragrams showing the behavioral phenotype of biased and non-biased animals considering the relative frequency of choice (tetragram frequency of choice*100/total number of turns).** Data were represented as mean ± S.E.M. and analyzed by linear mixed effects, followed by Tukey’s multiple comparison test. Asterisks indicates statistical differences compared to non-biased group or between biased groups (*p < 0.05, **p <0.01, ***p <0.001 and **\*\*\*\*p<0.0001**, n = 47 non-biased, n = 28 left-biased and n = 28 right-biased group).

### 2.2. Short-term avoidance memory and novelty response of biased animals

Although no interaction effect bias *vs.* shock (F_(2, 98)_ =1.259; *p*= 0.312) was observed for the Pavlovian responses, a significant effect for bias (F_(2, 98)_ =3.128; *p*= 0.035) and **conditioning (probe *vs* baseline)** (F _(1, 98)_ = 79.47; *p*<0001) effect was revealed. **ANOVA analysis are often underpowered and therefore unable to detect the significance of interaction terms** (Wahlsten 1990)**. Therefore post-hoc analysis was performed to specifically analyze the effects between groups. In general, all biased (*p<*0.0001 for right- and left-bias) and non-biased (*p<*0.0001) animals had a decreased time spent in the preference for conditioned stimulus (probe *vs* baseline)**. **However, both left- and right-biased animals (*p*<0.05) spent significantly less time in the conditioned area during the probe trial compared to non-biased group** (Fig. 3), **despite no significant differences in their baseline preferences.** No significant effect was observed for bias in all novel tank diving test-related parameters, including distance travelled (F _(2, 43)_ = 0.683; *p* = 0.510), immobility (F _(2, 43)_ = 2.348; *p* = 0.107), time in tank zones time (F _(2, 129)_ = 0.084; *p* = 0.918) (Fig. 4) and thigmotaxis (F _(2, 43)_ = 1.289; *p* = 0.286) (Fig. 5).

**Figure 3.**
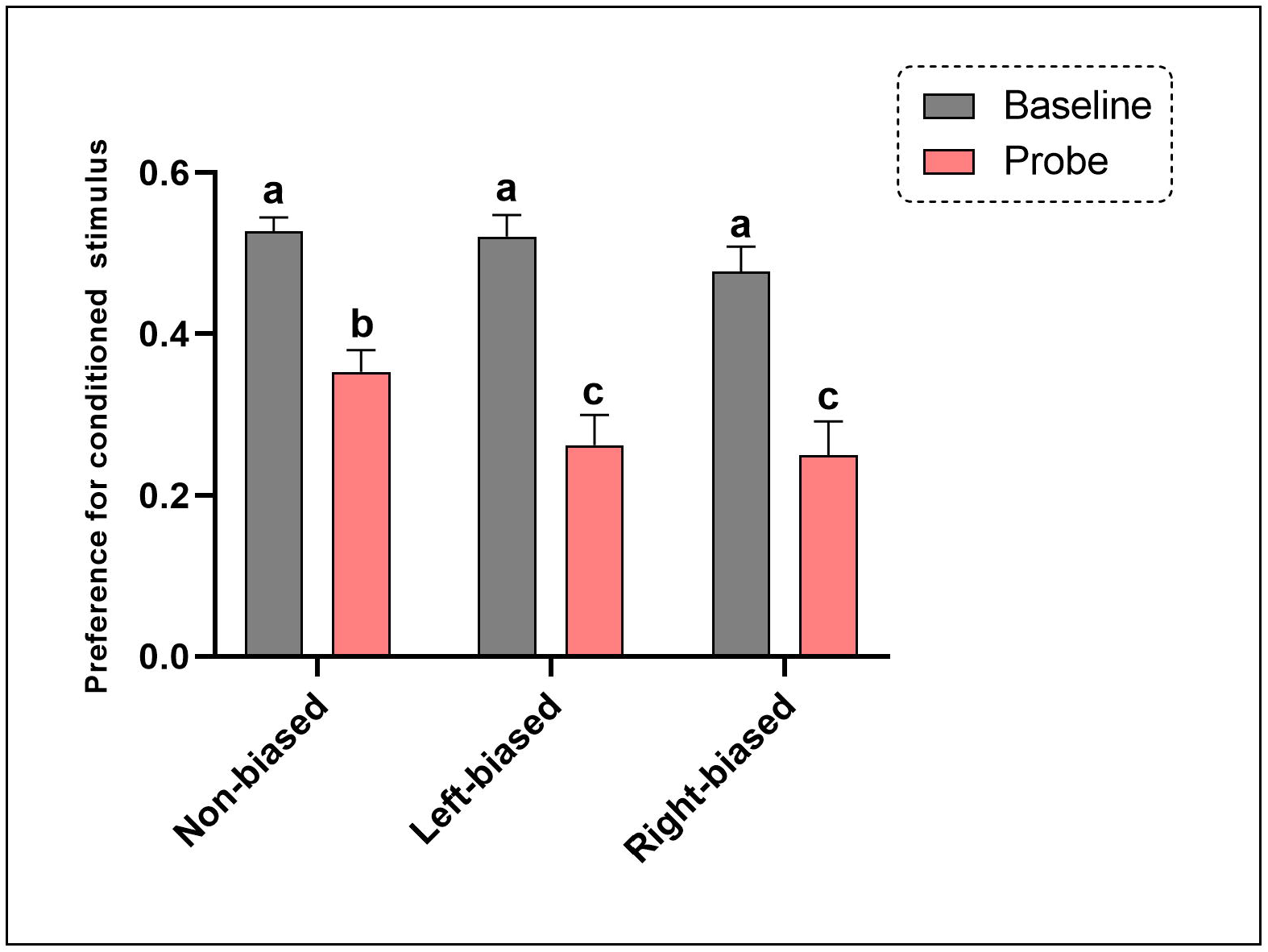
Left- and right-bias are related to fear avoidance learning responses in adult zebrafish. Data were represented as mean ± S.E.M. and analyzed by two-way RM ANOVA, followed by Tukey’s multiple comparison test. **Different letters indicate significant differences between groups (*p < 0.05*;** *n* = 25 non-biased, *n* = 17 left-biased and *n* = 13 right-biased group).

**Figure 4.**
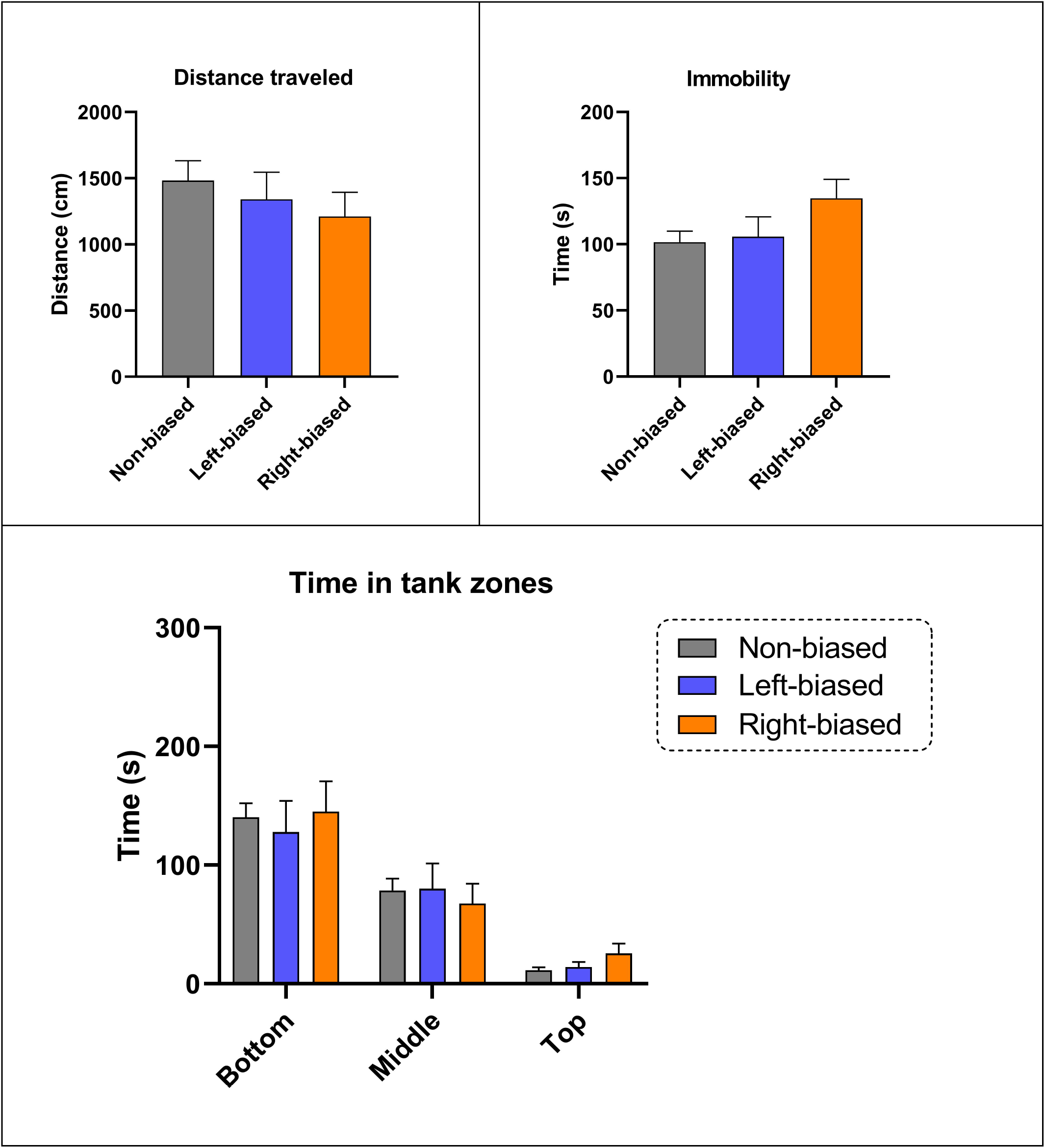
Behavioral laterality is not related to locomotor or anxiety-related phenotypes in adult zebrafish. Data were represented as mean ± S.E.M. and analyzed by **one-or two-way ANOVA** (*n* = 22 non-biased, *n* = 11 left-biased and *n* = 15 right-biased group).

**Figure 5.**
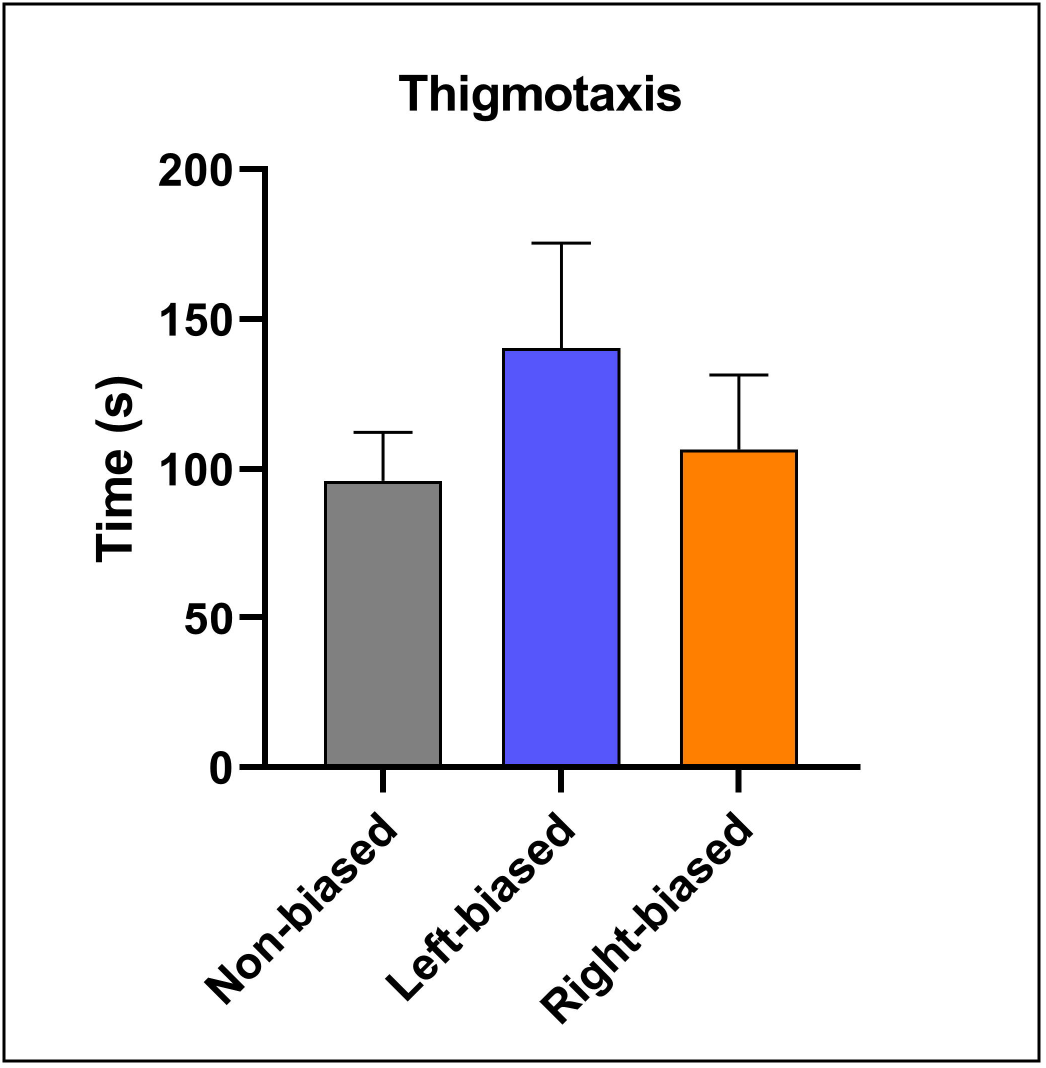
Left- and right-bias do not change thigmotaxis in adult zebrafish. Data were represented as mean ± S.E.M. and analyzed by **one-way ANOVA** (*n* = 20 non-biased, *n* = 11 left-biased and *n* = 15 right-biased group).

## 4. Discussion

In this study we evaluated **spontaneously occurring motor left-right bias** in adult zebrafish using **an** unconditioned Y-maze **protocol and** evaluated the predictive validity of **spontaneous behavioral laterality** on both unconditioned and conditioned measures of fear **and anxiety**. We showed, for the first time, that the zebrafish presents natural behavioral laterality in the Y-maze test **when analyzed using 16 overlapping tetragrams to evaluate exploration strategy**, suggesting that the protocol may be useful for screening this species for behavioral asymmetry. Second, we found that behaviorally lateralized animals show **a decrease in the relative number of alternation** responses and increase **in relative number of repetitions, in the Y-maze**. **This suggests that search strategies used by spontaneously biased individuals differs from that used by non-biased fish and is quite likely directly due to the lateralization.** Third, we observed that behavioral asymmetry predicts increased learning in a Pavlovian fear conditioning protocol but did not predict measures of unconditioned anxiety (novel tank test, thigmotaxis). Collectively, these data suggest, contrary to theories that laterality is related to increased stress-reactivity, that increased behavioral laterality may be related to increased cue-reactivity, particularly in relation to aversive cues. This has connotations for translational models of human disorders of affective state, in which heightened attention to threat-related cues is observed (Lichtenstein-Vidne et al. 2017) and in which brain and behavioral laterality are thought to be risk factors (Bruder et al. 2016).

**Left-right asymmetries in behavioral protocols including the T-maze and Y-maze have been widely utilized in rodents (Andrade et al. 2001; Nakagawa et al. 2004; Rodriguez et al. 1992). Here, for the first time, we observed that approximately a quarter of zebrafish present substantial natural left- (27.18 %) or right- (27.18%) locomotor lateralization, with the remaining 45.63%** of **animals showing stochastic patterns of left/right. These data are somewhat at odds from observed** bias in other models, in which there is a high number of right-biased animals (52.8%) and low numbers of left- (22.2%) and non-biased responses (25%) for rodents (Andrade et al. 2001). **A previous study (Chivers et al. 2016) showed that wild-caught fish, yellow-and-blueback fusiliers (*Caesio teres*), also present a natural increased bias for the right turn (65%) and decreased left bias (24%) and no biased (11%). However, they also observed that the strength of the behavioral lateralization was increased depending on predation pressure. Although the authors also did not find significant differences between the left- and right-biased groups, in general, the biased groups presented distinct patterns of behavior compared to non-biased animals. As mentioned earlier, larvae and adult zebrafish exhibit different patterns of left- and right eye use that correlate with boldness and response to novelty (Barth et al. 2005), aggression (Ariyomo and Watt 2013) and learning/memory (Andersson et al. 2015) behavior.**

**Studies focusing on the evolutionary aspects of lateralization suggest that behavioral asymmetries exist at population level because individually asymmetrical organisms coordinate their behavior with other asymmetrical organisms (Frasnelli 2013). The differences between left- and right-bias in other fish species. For example, right-biased animals showed increased escape responses (Chivers et al. 2016) and increased right-eye use when attacking the mirror image aggression (Bisazza and de Santi 2003). In invertebrates, laterality has been also consistent with the vertebrate findings, where eusocial ants show motor left-bias when exploring unfamiliar nest sites (Hunt et al. 2014) and eusocial honeybees have olfactory asymmetries that predict learning and recall of memory (Frasnelli et al. 2010; Rigosi et al. 2011; Rogers and Vallortigara 2008).**

**In general, brain and behavior lateralization are more common in social rather than in solitary species (Bisazza et al. 2000; Frasnelli 2013). Although zebrafish is a social species (Saverino and Gerlai 2008), research on zebrafish almost only uses domesticated strains, that were removed from wild populations many generations ago (Spence et al. 2008), as we used here. Zebrafish lab strains present genetic divergence from wild populations (Whiteley et al. 2011), and it is therefore important to consider that wild populations may have different lateralization patterns. Thus, behavioral patterns may change depending on stress-levels (Parker et al. 2012), predator pressure (Chivers et al. 2016), fish strain (Vignet et al. 2013) and animal origin (Spence et al. 2008), doing that more studies are needed to understand how behavioral laterality is influence by these variables in adult zebrafish.**

We observed that behavioral laterality has an important role in Y-maze performance, where left-right biased animals presented an increase of repetition behavior and decrease of alternation, in particular in the left-biased animals. **Previous studies by** (Cleal and Parker 2018; Gross et al. 2011) **and continuing work in our laboratory (data not shown here) in conjunction with rodent T- and Y-maze data suggest that the normal search strategy employed in mazes is to use a high level of alternations.** Alternations have been directly associated with functionally distinct search patterns, where the seeking for change and novelty may have a role in their exploratory profiles (Kool et al. 2010). **Rodriguez et al. (1992) described decreases in pure alternations and increases in pure repetition behavior in lateralized animals, and demonstrated that both stress and over-training decrease the alternation/repetition ratio through the promotion of an increase of biased responses. Although the dominant strategy, alternations are broken up with other exploratory patterns, as indicated in Fig 2B showing the relative frequency of each of the 16 tetragrams over the duration of a 1-hour trial. This discontinuous, but dominant search strategy, leads us to believe that the exploration patterns in the Y-maze are possibly associated with short term memory rather than motor memory (McBride and Parker 2015). However, further work would have to be carried out to determine to what extent, if any, alternations and repetitions are implicated in short term memory in the Y-maze. Here, we confirmed that both alternations and repetitions remain as a highly reliable behavioral pattern that is conserved across species (Ghafouri et al. 2016; Lewis et al. 2017; Pickering et al. 2015).**

In agreement with previous studies (Andrade et al. 2001), we showed that locomotor lateralization is associated with increased learning in a **Pavlovian** fear conditioning protocol. **Studies using other fish species showed that lateralized animals have a better response in cognitive tasks such as spatial reorientation (Sovrano et al. 2005), and left-eye bias is related to faster learning in conditioning tasks (Bibost and Brown 2014). In general**, the most predominant theory of how left-right bias affects learning and cognitive processing relates to a hypothesized increased stress-reactivity in lateralized animals (Carlson and Glick 1989; Neveu 1996; Westergaard et al. 2001). Interindividual differences in laterality have been shown to covary with, or predict, individual differences in stress-reactivity and susceptibility to stress-related pathology (Byrnes et al. 2016; Carlson and Glick 1989; Fride and Weinstock 1989; Ocklenburg et al. 2016). Here, we tested the hypothesis that left- and right-biased animals would differ in measures of stress-reactivity and anxiety-like phenotypes (Blaser and Rosemberg 2012; Egan et al. 2009; Parker et al. 2012). We found no significant differences in lateralized animals in our measures of anxiety, suggesting that the observed differences in behavioral phenotypes observed in the Y-maze and shock avoidance learning seems to not be related to stress-reactivity responses *per se*. Instead, our data may suggest that the lateralized fish are more reactive to stress-related cues. This would explain the increased performance **in** the Pavlovian fear conditioning, the fact that there were no differences in measures of anxiety, and the **altered search strategy in** the Y-maze test.

There are several theories regarding the mechanisms underlining behavior laterality in simple maze tasks. Diaz Palarea et al. (1987) were the first to report that left-right biased animals, as assessed via spatial asymmetry in a T-maze, had alterations in dopaminergic (DA) signaling. In addition, apomorphine (DA receptor agonist) and 6-hydroxydopamine lesions alters behavioral laterality of animals in the T-maze test (Castellano et al. 1987) and Y-maze (Nakagawa et al. 2004), confirming the involvement of DA system in behavioral asymmetry. DA receptors are strongly implicated in emotional learning and recall of emotionally relevant events in rats. For example, activation of D4-receptors in the medial pre-frontal cortex potentiates fear-associated memory formation but has no impact on recall (Lauzon et al. 2009; Laviolette et al. 2005), whereas activation of D1-like receptors blocks recalls of previously learned fear-associated memories but has no impact on learning (Lauzon et al. 2009), suggesting a double dissociation of function. Interestingly, the serotonergic (5-HT) system has also been shown to have an important role in mediating individual differences in anxiety-like responses and locomotor activity in zebrafish and exerts a minor modulatory role of the DA system (Tran et al. 2016). Both behavioral laterality and aversive memory is mostly associated with **modulatory action of the DA system**, but the 5-HT system has a major role modulating zebrafish responses to novelty. The precise mechanisms of how behavioral laterality modulates neuropsychiatric conditions are yet to be firmly established, and further studies are required to better understand the mechanisms in which behavioral laterality modulates aversive memory in zebrafish.

## 5. Conclusion

Overall, we showed for the first time that zebrafish exhibit **spontaneously occurring motor lateralization which can influence aversive learning responses. We found that biased animals show an altered search exploratory pattern Y-maze and increased** in learning in a Pavlovian fear conditioning protocol. Coupled with a lack of differences between lateralized and non-lateralized animals in unconditioned tests of anxiety, our data suggest that lateralized zebrafish may **show heightened** reactivity to fear related cues. These results have important connotations for translational models of depression and anxiety, particularly in the light of well-established links between laterality and anxiety/depression in humans. Finally, because biased animals present different behavioral performances in the Y-maze and Pavlovian fear conditioning protocols, left- and right-preference should be considered when working with zebrafish behavior, particularly to control variability in performance on more complex tasks.

## Compliance with Ethical Standards

### Funding

This study was financed in part by the Coordenação de Aperfeiçoamento de Pessoal de Nível Superior - Brazil (CAPES) - Finance Code 001 at the University of Portsmouth, UK. MOP receives funding from the Foundation for Liver Research and the British Academy. MC is supported by a Science Faculty Studentship from the University of Portsmouth.

### Ethical approval

All experiments were carried out following scrutiny by the University of Portsmouth Animal Welfare and Ethical Review Board, and under license from the UK Home Office (Animals (Scientific Procedures) Act, 1986) [PPL: P9D87106F].

